# Metabolic immaturity of newborns and breast milk bile acid metabolites are the central determinants of heightened neonatal vulnerability to norovirus diarrhea

**DOI:** 10.1101/2024.05.01.592031

**Authors:** Amy M. Peiper, Joyce Morales Aparicio, Lufuno Phophi, Zhengzheng Hu, Emily W. Helm, Matthew Phillips, Caroline G. Williams, Saravanan Subramanian, Michael Cross, Neha Iyer, Quyen Nguyen, Rachel Newsome, Christian Jobin, Stephanie N. Langel, Filemon Bucardo, Sylvia Becker-Dreps, Xiao-Di Tan, Paul A. Dawson, Stephanie M. Karst

## Abstract

Noroviruses are the leading global cause of acute gastroenteritis, responsible for 685 million annual cases. While all age groups are susceptible to noroviruses, children are vulnerable to more severe infections than adults, underscored by 200 million pediatric cases and up to 200,000 deaths in children annually. Understanding the basis for the increased vulnerability of young hosts is critical to developing effective treatments. The pathogenic outcome of any enteric virus infection is governed by a complex interplay between the virus, intestinal microbiota, and host immune factors. A central mediator in these complex relationships are host- and microbiota-derived metabolites. Noroviruses bind a specific class of metabolites, bile acids, which are produced by the host and then modified by commensal bacterial enzymes. Paradoxically, bile acids can have both proviral and antiviral roles during norovirus infections. Considering these opposing effects, the microbiota-regulated balance of the bile acid pool may be a key determinant of the pathogenic outcome of a norovirus infection. The bile acid pool in newborns is unique due to immaturity of host metabolic pathways and developing gut microbiota, which could underlie the vulnerability of these hosts to severe norovirus infections. Supporting this concept, we demonstrate herein that microbiota and their bile acid metabolites protect from severe norovirus diarrhea whereas host-derived bile acids promote disease. Remarkably, we also report that maternal bile acid metabolism determines neonatal susceptibility to norovirus diarrhea during breastfeeding by delivering proviral bile acids to the newborn. Finally, directed targeting of maternal and neonatal bile acid metabolism can protect the neonatal host from norovirus disease. Altogether, these data support the conclusion that metabolic immaturity in newborns and ingestion of proviral maternal metabolites in breast milk are the central determinants of heightened neonatal vulnerability to norovirus disease.

## INTRODUCTION

Human noroviruses are the leading global cause of childhood diarrhea and acute gastroenteritis in all age groups, attributed to a remarkable 685 million cases and the deaths of 50,000-200,000 young children each year^1–3^. Understanding their pathogenesis, particularly in young hosts, is thus an important priority to aid in the development of vaccines and therapeutics. Because noroviruses are enteric pathogens, a complete understanding of their pathogenesis requires characterization of viral interactions with intestinal microbiota encountered in the gut lumen and the consequences of these interactions. Gut microbiota comprise a dynamic ecosystem along the gastrointestinal tract, with trillions of microorganisms shaping host physiology, immune response, and pathogen susceptibility^4–6^. Over the past decade, it has become well-established that gut microbiota play an important role in regulating enteric virus infections^7–12^ and that microbiota-derived metabolites can mediate these effects^13^.

Noroviruses have been reported to bind a specific class of metabolites, bile acids^14–17^. Bile acids are synthesized in the liver from cholesterol where they are conjugated to either taurine or glycine, resulting in conjugated primary (host-derived) bile acids. These are stored in the gallbladder until feeding, when they are secreted into the duodenum. Once in the intestinal lumen, bacterial enzymes metabolize the primary conjugated bile acids into unconjugated primary and secondary (microbiota-derived) bile acids^18–20^. 95% of bile acids are reabsorbed across the intestinal epithelium and circulate back to the liver through a process termed enterohepatic circulation. The intestinal transport of bile acids is largely mediated by the apical sodium-dependent bile acid transporter (ASBT)^21^. Noroviruses can bind a broad range of bile acids, including host-derived and microbiota-biotransformed classes^14–17^. Interaction with host-derived bile acids has been shown to be proviral for both human and murine noroviruses in vitro through a range of mechanisms, including directly enhancing receptor binding and indirectly increasing the susceptibility of host cells to infection^14,22^. On the other hand, microbiota-derived bile acids can inhibit murine norovirus infection of adult mice in an interferon (IFN)-dependent manner^23^. Gut microbiota-biotransformed bile acids also protect from alphavirus, influenza virus and coronavirus infections^24,25^, suggesting that this antiviral process functions across virus families and at both local and systemic tissues. Altogether, these findings demonstrate that host-derived bile acids promote norovirus infection while microbiota-derived bile acids activate an antiviral immune response and inhibit infection, leading to the model that the microbiota-regulated balance of bile acid species present in the gut lumen determines norovirus susceptibility.

Intriguingly, the bile acid pool in neonates is markedly distinct from adults for multiple reasons^26–28^. First, genes involved in bile acid synthesis and hepatobiliary transport are age-regulated in humans and rodents, increasing steadily over the first month of life in mice^26–28^. Second, the abundance of commensal bacteria expressing enzymes necessary for conversion of host-derived bile acids to microbiota-dependent species is low at birth and steadily increases through weaning^27,28^. Third, enterohepatic circulation is prevented in neonates by postnatal repression of the main intestinal bile acid transporter ASBT^21,29^. Finally, breastfed neonates receive maternal bile acids in breast milk that contribute to their bile acid pool^30,31^. The net effect of these developmental factors is that the neonatal bile acid pool could be heavily dominated by host-derived bile acids. Considering the potential proviral activity of these bile acid species, it is possible that age-specific homeostatic bile acid metabolism is the central determinant of the severity of a norovirus infection and that directed targeting of this pathway could prevent norovirus infections. Indeed, herein we report that host-derived and microbiota-derived bile acids have opposing effects in regulating murine norovirus (MNV) disease severity, playing exacerbating and protective roles, respectively. Remarkably, maternal bile acids delivered to the neonate in breast milk govern neonatal vulnerability to MNV diarrhea and microbiota maturation can provide protection by biotransforming the bile acid pool.

## RESULTS

### Microbiota-mediated bile acid metabolism protects neonates from severe MNV diarrhea

While adult mice are resistant to MNV-induced disease, neonatal mice infected with MNV develop acute self-resolving diarrhea in a time course that mirrors human norovirus disease^32–34^. This model system enabled us to test the role of gut microbiota in regulating norovirus diarrhea. Based on published methods^12^, neonatal mice were given two doses of ABX by intragastric (i.g.) inoculation (1 d before infection and 1 d after infection) which was sufficient to fully deplete intestinal microbiota. Importantly, we selected a cocktail of the three most commonly prescribed ABX in human neonates^35^. ABX-treated pups were infected with 10^7^ TCID_50_ units of the diarrheagenic MNV strain WU23^33^. Microbiota depletion significantly exacerbated MNV-induced diarrhea, as indicated by increased disease severity and incidence **(Fig. 1a)**. A similar phenotype was observed in neonates infected at three days of age (postnatal 3 [P3]) or P5. Another indicator of intestinal disease is increased permeability of the epithelial barrier which can be assessed by measuring the amount of orally administered fluorescently labeled dextran that reaches circulation. ABX treatment of neonatal mice resulted in significantly increased intestinal permeability during MNV infection **(Fig. 1b)**. There was a positive correlation between fecal scores and levels of systemic FITC-Dextran 10 kDa (FD10) **(Supplementary Fig. 1)**. Certain antibiotics can have microbiota-independent antiviral effects^36–40^. To test whether ABX administration exacerbated WU23-induced diarrhea in a microbiota-dependent or microbiota-independent manner, we compared diarrhea severity in germ-free mice treated with PBS or ABX. Consistent with microbiota-dependent activity, ABX treatment had no effect on diarrhea severity in germ-free pups **(Fig. 1c)**. We have previously reported strict age restriction of MNV-induced diarrhea, with pups P5 or younger experiencing diarrhea but minimal disease observed in mice infected P7 or older^32,33^. Microbiota depletion extended the window of susceptibility since P7 and P10 ABX-treated mice developed diarrhea of comparable severity and incidence as P5 pups **(Fig. 1d)**. Thus, intestinal microbiota provide protection from severe WU23-induced diarrhea. Our prior studies in adult mice revealed that microbiota-mediated effects on acute MNV infection are largely a consequence of microbial regulation of the bile acid pool, with microbiota-derived unconjugated bile acids inhibiting duodenal infection^23^. To determine whether microbiota-mediated metabolism of bile acids was likewise responsible for protecting from severe disease, ABX-treated neonates were reconstituted with *Clostridium scindens*, a bacterial species with potent bile acid biotransforming activity^41,42^. ABX-treated, *C. scindens*-reconstituted pups were fully protected from WU23-induced diarrhea in contrast to control ABX-treated pups receiving bacterial growth media **(Fig. 1e)**. Gut microbiota modify bile acids in numerous ways but the gateway modification is removal of the taurine or glycine residue added during the host conjugation process. To determine whether this deconjugation step is required for *C. scindens*-mediated protection, ABX-treated pups were administered caffeic acid phenethyl ester (CAPE), a potent inhibitor of the bacterial bile salt hydrolase (BSH) gene responsible for bile acid deconjugation^43^, prior to *C. scindens* reconstitution. Confirming a role for bacterial deconjugation, CAPE pretreatment abolished the protective effect of *C. scindens* reconstitution **(Fig. 1f)**.

**Figure 1.**
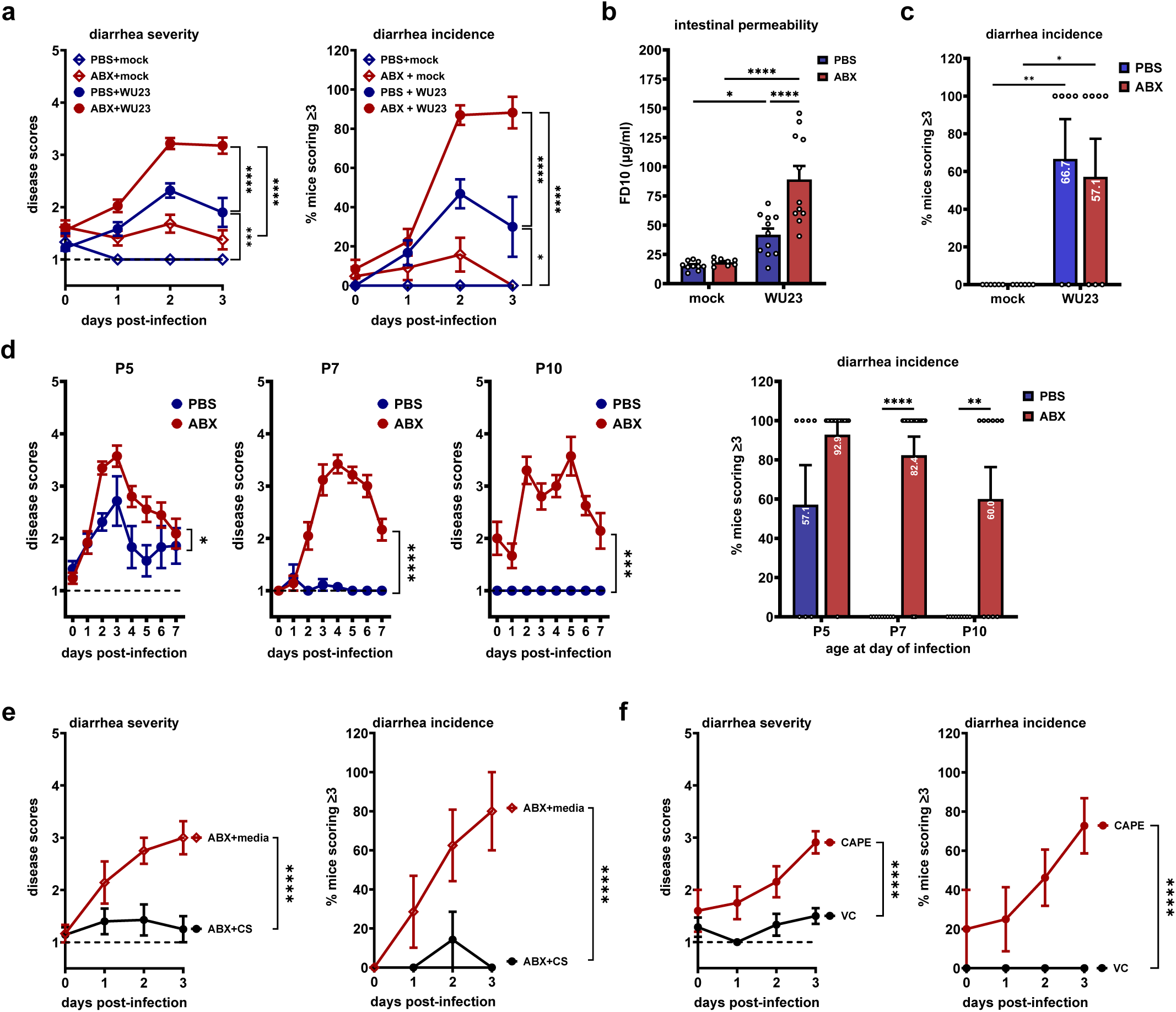
Microbiota-mediated bile acid metabolism protects neonates from severe MNV diarrhea. **a)** Neonatal B6 mice were treated with PBS or ABX via i.g. injection at P2, infected i.g. with 1 x 10^7^ TCID_50_ units of WU23 or mock inoculum at P3, and treated with PBS or ABX again at P4. Mice were monitored for disease severity and incidence for 3 dpi. Due to the nature of the fecal scoring system described in the methods, group sizes varied by time point (PBS+WU23: *n* = 27 [0 dpi], 36 [1 dpi], 47 [2 dpi], 10 [3 dpi]; ABX+WU23: *n* = 36 [0 dpi], 40 [1 dpi], 46 [2 dpi], and 17 [3 dpi]; PBS+mock: *n* = 9 [0 dpi], 9 [1 dpi], 7 [2 dpi] 7 [3 dpi]; and ABX+mock: *n* = 21 [0 dpi], 22 [1 dpi], 19 [2 dpi], 8 [3 dpi]). **b)** To measure intestinal permeability, PBS- or ABX-treated neonates infected i.g. with 1 x 10^7^ TCID_50_ units of WU23 (PBS: *n* = 11; ABX: *n* = 11) or mock inoculum (PBS: *n* = 10; ABX: *n* = 8) were administered 40 mg/kg FD10 via i.g. injection at 69 hpi and the level of FD10 in serum measured at 72 hpi. **c)** Germ-free B6 neonates were treated with PBS or ABX at P2 and P4 and infected with 1 x 10^7^ TCID_50_ units of WU23 (PBS: *n* = 6; ABX: *n* = 7) or mock inoculum (PBS: *n* = 6; ABX: *n* = 6) at P3. Disease incidence was measured at 2 dpi. **d)** Neonatal B6 mice administered PBS or ABX 1 d before and 1 d after infection were infected with 1 x 10^7^ TCID_50_ units of WU23 on P5 (PBS: *n* = 12 [0 dpi], 20 [1 dpi], 31 [2 dpi], 7 [3 dpi], 6 [4 dpi], 7 [5 dpi], 6 [6 dpi], 7 [7 dpi]; ABX: *n* = 17 [0 dpi], 25 [1 dpi], 29 [2 dpi], 14 [3 dpi], 11 [4 dpi], 9 [5 dpi], 9 [6 dpi], 11 [7 dpi]), P7 (PBS: *n* = 6 [0 dpi], 4 [1 dpi], 7 [2 dpi], 9 [3 dpi], 14 [4 dpi], 9 [5 dpi], 4 [6 dpi], 3 [7 dpi]; ABX: *n* = 11 [0 dpi], 14 [1 dpi], 21 [2 dpi], 17 [3 dpi], 19 [4 dpi], 14 [5 dpi], 10 [6 dpi], 12 [7 dpi]), or P10 (PBS: *n* = 8 [0 dpi], 7 [1 dpi], 9 [2 dpi], 9 [3 dpi], 6 [4 dpi], 5 [5 dpi], 5 [6 dpi], 5 [7 dpi]; ABX: *n* = 5 [0 dpi], 9 [1 dpi], 10 [2 dpi], 10 [3 dpi], 10 [4 dpi], 7 [5 dpi], 8 [6 dpi], 7 [7 dpi]). Mice were monitored for disease severity and incidence up to 7 dpi. **e)** B6 neonates were administered ABX at P2, reconstituted with 1 x 10^6^ CFU of *C. scindens* (*n* = 7 [0 dpi], 5 [1 dpi], 7 [2 dpi], 4 [3 dpi]) or bacterial media (*n* = 6 [0 dpi], 7 [1 dpi], 8 [2 dpi], 5 [3 dpi]) at P3, and infected with 1 x 10^7^ TCID_50_ units of WU23 at P5. They were monitored for disease severity and incidence for 3 dpi. **f)** B6 neonates were administered ABX at P2, reconstituted with 1 x 10^6^ CFU of *C. scindens* at P3, treated with 75mg/kg of CAPE (*n* = 5 [0 dpi], 8 [0 dpi], 13 [0 dpi], 11 [0 dpi]) or vehicle control (*n* = 7 [0 dpi], 7 [0 dpi], 6 [0 dpi], 12 [0 dpi]) at P4 and P6, and infected with 1×10^7^ TCID_50_ units of WU23 at P5. Mice were monitored for disease incidence and severity for 3 dpi. Statistical significance was calculated using 2-way ANOVA with Tukey’s multiple comparison test for all panels. *Due to the nature of our fecal scoring system which requires that a pup defecate upon stomach palpation at each time point in order to generate material to be scored, group sizes differ by time point. We have included precise group sizes per time point per condition to align with journal specifications but, understanding that this is cumbersome to read, we used gray text to help with readability.

### Activation of the bile acid receptor TGR5 protects from MNV disease

We next sought to directly test the role of microbiota-derived bile acids in protecting from MNV-induced disease. Supplementation of ABX-treated neonatal mice with an unconjugated secondary bile acid deoxycholic acid (DCA) fully rescued the protection afforded by gut microbiota, as indicated by reduced diarrhea severity, disease incidence, and intestinal permeability compared to control mice treated with vehicle control **(Fig. 2a)**. To understand how microbiota-derived bile acids inhibit MNV infection as this knowledge could guide the design of rational antivirals, it is important to note that they act as signaling molecules, interacting with bile acid-specific receptors expressed by many cell types^44^. Engagement of bile acid receptors activates a negative feedback loop to limit production of bile acids while also regulating inflammatory pathways^45^. Moreover, activation of the G protein-coupled bile acid receptor GPBAR1 (TGR5) inhibits replication of numerous viruses^25,46–48^. Based on its broad antiviral activity, we questioned whether it likewise functions in an antiviral manner during WU23 infection. Indeed, pretreatment of neonatal mice with the TGR5 agonist INT-777 (semi-synthetic unconjugated cholic acid [CA] analogue^49^) for 3 days prior to infection and 1 dpi provided protection from WU23-induced disease in a dose-dependent manner, as indicated by reduced diarrhea severity, incidence, and intestinal permeability **(Fig. 2b)**. This activity was dependent on TGR5 since INT-777 displayed no protective effect in *Tgr5^-/-^* mice **(Fig. 2c)**. Consistent with a role for TGR5-dependent amplification of type I interferon (IFN) production as has been reported previously for other viruses^46^, IFN-β induction was accelerated in the small intestines of WU23-infected mice pretreated with INT-777 compared to control mice **(Fig. 2d)**. Altogether, these results support the conclusion that microbiota-derived bile acids can bind TGR5 and drive an antiviral response to prevent WU23 disease. Yet it is important to note that vehicle control-treated *Tgr5^-/-^* neonates developed diarrhea of comparable severity and incidence to their wild-type counterparts **(Fig. 2c)**, implying that endogenous TGR5 activation does not occur robustly enough in neonates to provide protection. This could be explained by low abundance of metabolized bile acids early in life, consistent with a paucity of bacterial species capable of metabolizing bile acids in the early-life microbiota^27,50^ and decreased enterohepatic circulation of bile acids early in life to prevent toxic accumulation of bile acids in hepatocytes and ileal enterocytes^51,52^.

**Figure 2.**
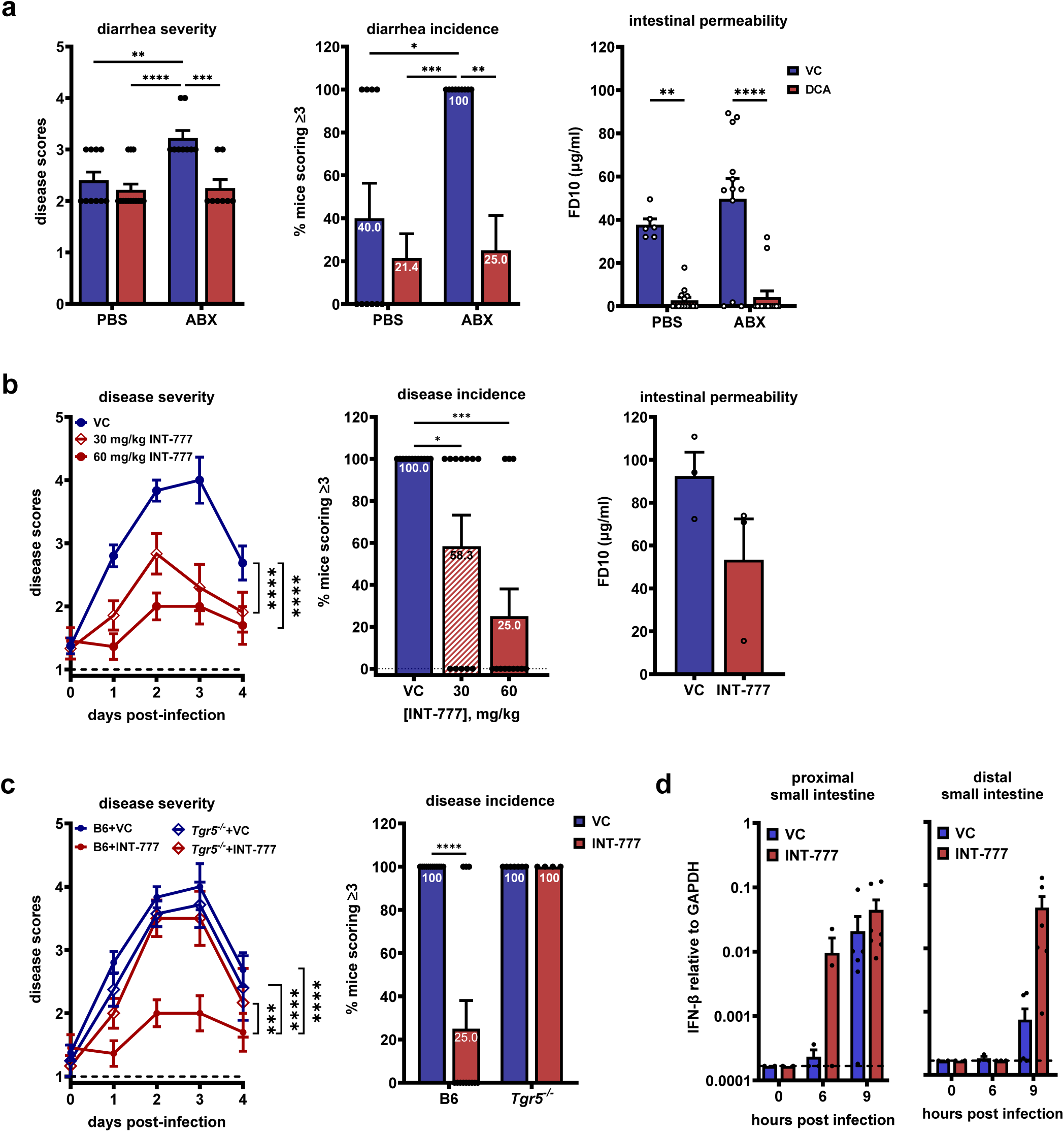
Activation of the bile acid receptor TGR5 protects from MNV disease. **a)** B6 neonatal mice were treated with PBS or ABX at P2, P4, and P6, administered 70 µg/g of DCA (PBS: *n* = 14; ABX: *n* = 8) or vehicle control (VC; PBS: *n* = 10; ABX: *n*= 9) at P3, P4, and P5 via i.g. injection, and infected with 1 x 10^7^ TCID_50_ units of WU23 at P5.5. Disease severity and incidence were determined at 2 dpi. Separate groups of mice treated in the same manner were used to assess intestinal permeability by administering 40 mg/kg FD10 at 45 hpi and measuring FD10 in serum at 48 hpi (PBS+VC: *n* = 6; ABX+VC: *n* = 12; PBS+DCA: *n* = 14; ABX+DCA: *n* = 14). **b)** Neonatal B6 mice were treated with VC (*n* = 16 [0 dpi], 15 [1 dpi], 12 [2 dpi], 6 [3 dpi], 16 [4 dpi]), 30 mg/kg INT-777(*n* = 9 [0 dpi], 14 [1 dpi], 12 [2 dpi], 10 [3 dpi], 11 [4 dpi]) or 60 mg/kg INT-777 (n= 11 [0 dpi], 11 [1 dpi], 12 [2 dpi], 13 [3 dpi], 10 [4 dpi]) at P2, P3, P4 and P6 via i.g. injection, and infected at P5 with 5 x 10^7^ TCID_50_ units of WU23. Mice were monitored for disease severity and incidence for 4 dpi. The incidence at 2 dpi is shown. Pups treated in the same manner were used to assess intestinal permeability (*n* = 3 pooled litters per group). **c)** P3 B6 or *Tgr5^-/-^* neonatal mice were treated with VC (B6: *n* = 16 [0 dpi], 15 [1 dpi], 12 [2 dpi], 6 [3 dpi], 16 [4 dpi]; *Tgr5^-/^*:*^-^ n* = 4 [0 dpi], 8 [1 dpi], 7 [2 dpi], 7 [3 dpi], 5 [4 dpi]) or 60 mg/kg of INT-777 (B6: *n* = 11 [0 dpi], 11 [1 dpi], 12 [2 dpi], 13 [3 dpi], 10 [4 dpi]; *Tgr5^-/-^*: *n* = 6 [0 dpi], 9 [1 dpi], 4 [2 dpi], 6 [3 dpi], 6 [4 dpi]) at P2, P3, P4, and P6 and infected with 5 x 10^7^ TCID_50_ units of WU23 at P5. Mice were monitored for disease severity and incidence for 4 dpi. The incidence at 2 dpi is shown. **d)** B6 neonatal mice were treated with VC or 60 mg/kg of INT-777 at P2, P3 and P4 and infected with 5 x 10^7^ TCID_50_ units of WU23 at P5. Portions of the proximal and distal small intestines were harvested at 0 hpi (VC: *n* = 2, INT-777: *n* = 2), 6 hpi (VC: *n* = 3, INT-777: *n* = 3), or 9 hpi (VC: *n* = 6, INT-777: *n* = 6). *IFN-β* levels were determined using quantitative RT-PCR and normalized to the *GAPDH* housekeeping gene. Statistical significance was calculated using 2-way ANOVA with Tukey’s multiple comparison test for all panels.

### Host-derived bile acids exacerbate MNV disease

To begin testing the concept that differences in the bile acid pool underlie age-dependent susceptibility to norovirus disease, bile acid pools in neonatal and adult mouse feces were evaluated by quantifying 70 bile acid species (51 standard bile acids and 19 bile acid detoxification products) using ultra-high performance liquid chromatography-mass spectrometry (UPLC-MS; **Fig. 3a and Supplemental Data File 1)**. As expected, neonates excreted less total bile acids than adult mice **(Fig. 3b)**. Adult mice excreted predominantly unconjugated secondary (microbiota-metabolized) bile acids whereas neonates excreted nearly exclusively conjugated primary (host-derived) bile acids **(Fig. 3c)**. The abundance of five bile acid species was significantly different in neonatal and adult samples: two conjugated bile acids (taurocholic acid [TCA] and T-allo-CA) were more abundant in neonatal samples and three unconjugated secondary bile acids (DCA, ω-muricholic acid [ωMCA], and 12-keto chenodeoxycholic acid [12-keto CDCA]) were more abundant in adult samples **(Fig. 3d)**. In contrast to the protective role played by microbiota-derived bile acids like DCA, host-derived conjugated primary bile acids have been reported by several groups to promote norovirus infection in vitro^14,22^. To test whether they similarly promote pathogenesis in vivo, we investigated the effect of supplementing pups with TCA, the most abundant bile acid in neonatal feces. Consistent with a proviral role for host-derived bile acids in vivo, TCA supplementation exacerbated WU23-induced disease under bacteria-replete and bacteria-deplete conditions, as indicated by increased diarrhea severity and disease incidence **(Fig. 3e)** and increased intestinal permeability **(Fig. 3f)**. Another conjugated primary bile acid, glycochenodeoxycholic acid (GCDCA), has been reported to promote human and murine norovirus infections in vitro^14,15,22^ and it likewise promoted pathogenesis in neonatal mice comparably to TCA **(Fig. 3e)**. Altogether, these results demonstrate that host-derived bile acids exacerbate WU23-induced disease and they are the dominant bile acid class in neonatal hosts, supporting the possibility that heightened neonatal vulnerability to noroviruses derives from their metabolic immaturity.

**Figure 3.**
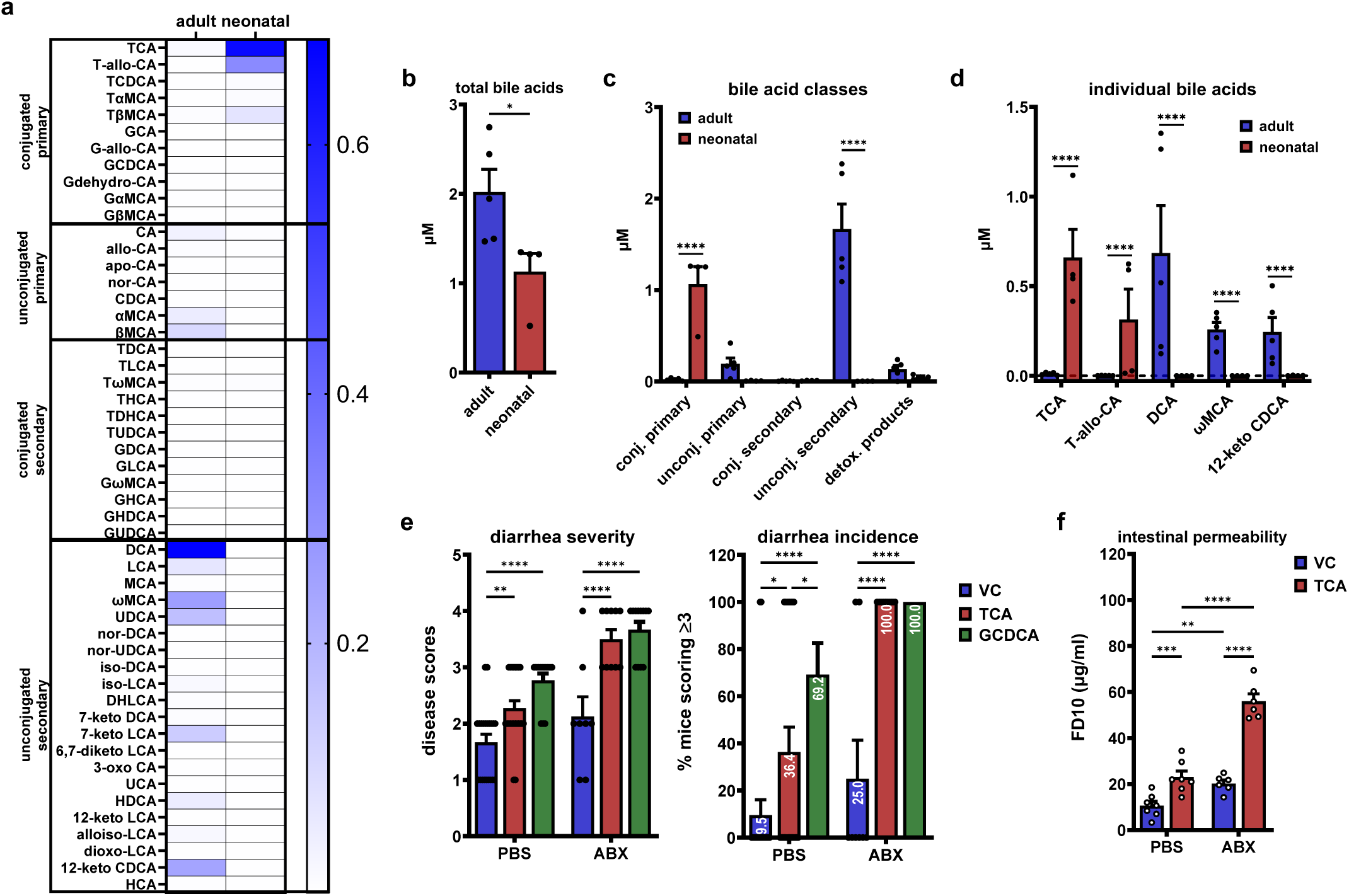
Host-derived bile acids exacerbate MNV disease. **a-d)** Fecal samples were collected from individual B6 adult mice (*n* = 5) or pooled P4 B6 litters (*n* = 4) and bile acid profiling was performed with UPLC-MS. A heatmap showing the mean concentrations of individual bile acids **(a)**, the abundance of total bile acid pools **(b)**, the abundance of bile acid categories **(c)**, and the abundance of individual bile acids that were statistically different between groups **(d)** are shown. **e)** B6 neonatal mice were treated with PBS or ABX at P2, P4, and P6, administered 70 µg/g of TCA (PBS: *n* = 22; ABX: *n* = 10), GCDCA (PBS: *n* = 13; ABX: *n* = 12), or VC (PBS: *n* = 21; ABX: *n* = 8) at P3, P4, and P5 via i.g. injection, and infected with 1 x 10^6^ TCID_50_ units of WU23 at P5.5. Disease severity and incidence were determined at 2 dpi. **f)** Separate groups of mice treated in the same manner were used to assess intestinal permeability at 48 hpi (PBS+VC: *n* = 7; ABX+VC: *n* = 6; PBS+TCA: *n* = 7; ABX+TCA: *n* = 6). Statistical significance was calculated using an unpaired student t-test **(b)** and 2-way ANOVA with Tukey’s multiple comparison test **(c-f)**.

### The main intestinal bile acid transporter ASBT determines breast milk bile acid pools and neonatal susceptibility to MNV

While the dominance of host-derived bile acids in neonates can be partly explained by immature microbiota early in life^26–29^, another contributing factor in breastfed neonates may be the nature of the breast milk bile acid pool. Forsyth et al. established the presence of bile acids in breast milk in 1983^30^ but the types of bile acids in breast milk have not been well-characterized. Blazquez et al. performed the most detailed analysis to date, identifying TCA as the most abundant of 21 species measured in milk collected from lactating mice^31^. To more comprehensively characterize the breast milk bile acid pool, we collected breast milk from lactating B6 dams and quantified 50 bile acid species using UPLC-MS **(Fig. 4a and Supplemental Data File 1)**. Consistent with Blazquez et al., TCA was the most abundant bile acid although there were other species including microbiota-metabolized (e.g., unconjugated and secondary) bile acids present **(Fig. 4a)**. In order to test whether breast milk bile acids influence neonatal susceptibility to norovirus disease, we sought a model system in which this bile acid pool was altered. Bile acids undergo enterohepatic circulation by trafficking from their site of synthesis in the liver through the small intestinal lumen to be reabsorbed across the intestinal epithelium and returned to the liver via portal circulation. Nutrients, microbiota, metabolites, and immune cells have been demonstrated to traffic from the maternal intestinal tract to the mammary gland during lactation, i.e. the entero-mammary pathway^53^. Since there were microbiota-dependent bile acids in the breast milk of wild-type dams, we reasoned that their most likely mode of trafficking to the mammary gland was following reabsorption across the gut mucosa. Apical sodium-dependent bile acid transporter (ASBT) is the main intestinal transporter of bile acids so we analyzed the breast milk bile acid pool in *Asbt^-/-^* dams **(Fig. 4a and Supplemental Data File 1)**. The composition of the pool was distinct from the pool in wild-type mice both quantitatively and qualitatively: There was a nearly 50% reduction in total bile acids in breast milk collected from *Asbt^-/-^* dams **(Fig. 4b)** due to a significantly reduced amount of conjugated primary bile acids **(Fig. 4c)**. There was significantly less TCA, TαMCA, TβMCA, and TωMCA, and a significant elevation of DCA **(Fig. 4d)**. To next explore the consequences of an altered breast milk bile acid pool on neonatal susceptibility to MNV diarrhea, P3 *Asbt^-/-^* or wild-type B6 control mice were infected with WU23 and monitored for diarrhea. Importantly, intestinal ASBT expression is suppressed in neonates by destabilization of the transcript^54–56^ so any difference in disease should result from maternal effects of ASBT deficiency. Remarkably, *Asbt^-/-^* neonates were nearly fully protected from WU23-induced diarrhea in contrast to B6 controls which developed diarrhea that peaked at 3 dpi and resolved by 4 dpi **(Fig. 4e)**. Protection was also evident when measuring intestinal permeability **(Fig. 4f)**. There are conditions that prematurely induce ASBT, including formula feeding^57^, glucocorticoid treatment^58^, pathophysiological conditions like necrotizing enterocolitis^59,60^, and deficiency of the bile acid transporter OSTα/β expressed on the basolateral surface of IECs^51^. Thus, it was theoretically possible that WU23 infection prematurely induces ASBT expression in neonatal mice in a manner that promotes viral infection. Because of the surprising nature of our results suggesting that maternal ASBT genotype determines neonatal susceptibility to WU23, we wanted to fully explore this alternative possibility. Using quantitative RT-PCR with ASBT-specific primers **(Supplementary Fig. 2)** and RNAscope-based in situ hybridization with an ASBT-specific probe **(Fig. 4g)**, we detected abundant ASBT expression in adult ileum but did not detect appreciable ASBT expression in mock-inoculated or WU23-infected neonates. These results rule out the possibility that viral induction of premature ASBT expression explains the striking phenotype observed in *Asbt^-/-^* pups.

**Figure 4.**
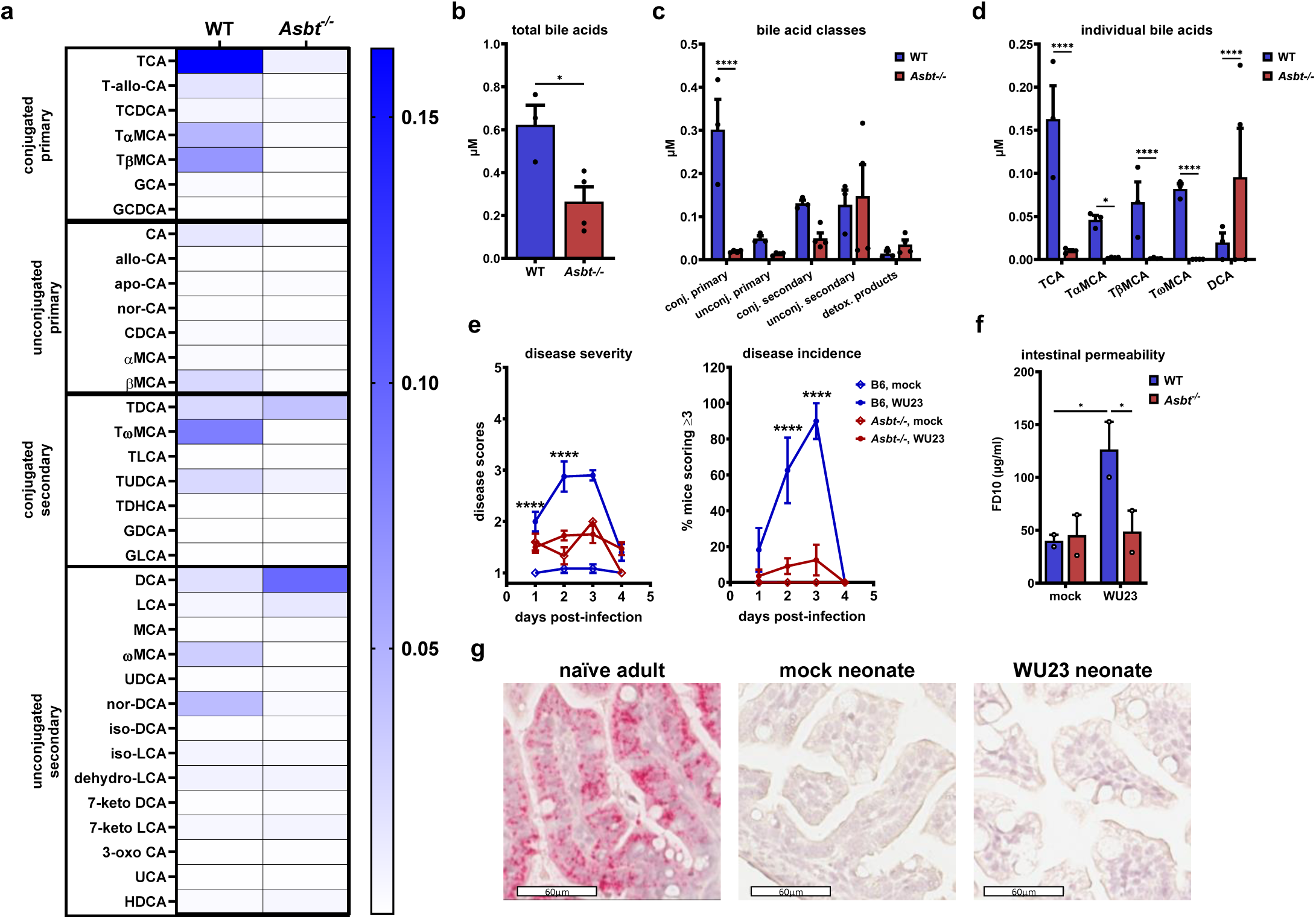
The main intestinal bile acid transporter ASBT determines breast milk bile acid pools and neonatal susceptibility to MNV. **a-d)** Breast milk samples were collected daily from naïve B6 or *Asbt^-/-^* lactating dams (*n* = 4) for 4 consecutive days and bile acid profiling was performed with UPLC-MS. A heatmap showing the mean concentrations of individual bile acids **(a)**, the abundance of total bile acid pools **(b)**, the abundance of bile acid categories **(c)**, and the abundance of individual bile acids that were statistically different between groups **(d)** are shown. **e)** B6 or *Asbt^-/-^* P3 mice were infected with 1 x 10^7^ TCID_50_ units of WU23 (B6: *n* = 11 [0 dpi], 8 [1 dpi], 10 [2 dpi], 11 [3 dpi]; *Asbt^-/^*:*^-^ n* = 28 [0 dpi], 44 [1 dpi], 16 [2 dpi], 17 [3 dpi]) or mock inoculum (B6: *n* = 11 [0 dpi], 12 [1 dpi], 12 [2 dpi], 10 [3 dpi]; *Asbt^-/-^*: *n* = 10 [0 dpi], 9 [1 dpi], 1 [2 dpi], 2 [3 dpi]). Mice were monitored for disease severity and incidence for 4 dpi. **f)** To measure intestinal permeability, B6 or *Asbt^-/-^* P3 mice were infected with 1 x 10^7^ TCID_50_ units of WU23 or mock inoculum, 40 mg/kg FD10 was administered at 45 hpi, and the level of FD10 in serum measured at 48 hpi. Serum was pooled from whole litters for analysis (*n* = 2 pooled litters per group). Statistical significance was calculated using an unpaired t-test **(b)** and 2-way ANOVA with Tukey’s multiple comparison test **(c-f)**. **g)** Intestinal tissue sections were collected from naïve adult B6 mice (*n* = 3) and from neonatal B6 mice infected with 1 x 10^8^ TCID_50_ units of WU23 or mock inoculum at 18 hpi (*n* = 5 per group) and hybridized with an ASBT-specific RNAscope probe. Representative images are shown.

### Maternal bile acids delivered via breast milk promote MNV diarrhea in the neonatal host

To more rigorously test the possibility that maternal ASBT expression determines neonatal susceptibility to WU23, fostering studies were performed in which *Asbt^-/-^* pups were fostered by wild-type dams so the breast milk consumed by pups had a normal bile acid composition but the pups themselves were ASBT-deficient. 64% of these pups developed diarrhea comparable to 63% of wild-type pups fostered with a wild-type dam **(Fig. 5a)**. Conversely, only 14% of wild-type pups and 6% of *Asbt^-/-^* pups fostered by *Asbt^-/-^* dams developed WU23-induced diarrhea **(Fig. 5a)**. Furthermore, TCA supplementation of *Asbt^-/-^* pups was sufficient to promote WU23 pathogenesis **(Fig. 5b)**, confirming that reduced delivery of host-derived bile acid species in the breast milk of *Asbt^-/-^* dams is responsible for protecting neonates from disease. In a complementary approach, wild-type B6 dams were treated with cholestyramine, a bile acid sequestrant, to deplete the bile acid pool available for enterohepatic circulation. Under these conditions, the severity and incidence of diarrhea in neonates inversely correlated with the concentration of cholestyramine fed to the dams **(Fig. 5c)**. Altogether, these findings reveal that maternal bile acids, specifically conjugated primary bile acids, delivered to neonates in breast milk increase MNV disease. Conjugated primary bile acids have been demonstrated to enhance human norovirus infection in vitro^22^ and infants are susceptible to severe human norovirus infections^61–63^, raising the possibility that breast milk bile acids can likewise exacerbate human norovirus disease. Consistent with this, approximately half of the bile acid pool in human breast milk is dominated by conjugated primary species as it is in mouse breast milk **(Fig. 5d and Supplemental Data File 1)**.

**Figure 5.**
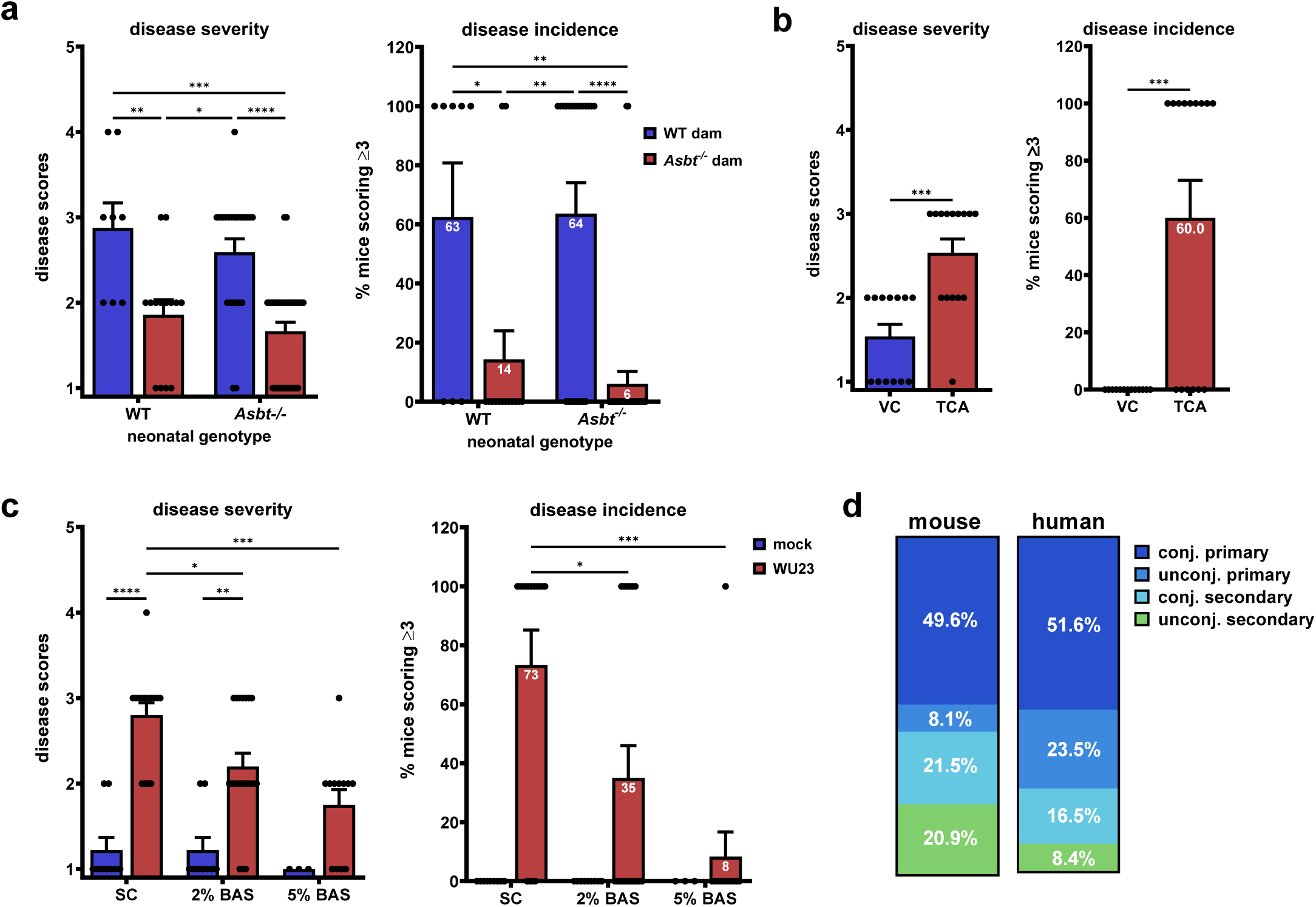
Maternal bile acids delivered via breast milk promote MNV diarrhea in the neonatal host. **a)** B6 or *Asbt^-/-^* neonatal mice fostered by dams of the same or opposite genotype at birth (B6 pups fostered by a B6 dam: *n* = 8; B6 pups fostered by an *Asbt^-/-^* dam: *n* = 22; *Asbt^-/-^* pups fostered by an *Asbt^-/-^* dam: *n* = 33; and *Asbt^-/-^* pups fostered by a B6 dam: *n* = 14) were infected with 1 x 10^7^ TCID_50_ units of WU23 at P3. Disease severity and incidence were determined at 2 dpi. **b)** *Asbt^-/-^* neonates were administered 70 µg/g of TCA (*n* = 15) or vehicle control (VC; *n* = 13) at P3, P4, and P5 and infected with 1×10^7^ TCID_50_ units of WU23 at P5.5. Disease severity and incidence were determined at 2 dpi. **c)** B6 dams were fed either standard chow (SC) or chow supplemented with 2% or 5% of the bile acid sequestrant (BAS) cholestyramine beginning the day she gave birth. Neonates from these dams were infected with 1 x 10^7^ TCID_50_ units of WU23 (SC: *n* = 15; 2% BAS: *n* = 20; 5% BAS: *n* = 12) or mock inoculum (SC: *n* = 9; 2% BAS: *n* = 9; 5% BAS: *n* = 5) at P5. Disease severity and incidence were determined at 2 dpi. **d)** The proportion of major bile acid classes in the total bile acid pool was compared between naïve B6 mouse breast milk (*n* = 3 pooled) and human breast milk (*n* = 24) samples. Statistical significance was calculated using a 2-way ANOVA with Tukey’s multiple comparison test (panels a, c) and unpaired student t-tests (panel b).

## DISCUSSION

Neonates are highly susceptible to many pathogens, a vulnerability that has long been assumed to be largely driven by immaturity of the neonatal immune system. Yet our studies support the concept that metabolic, not immunologic, immaturity underlies neonatal vulnerability to severe norovirus diarrhea. Specifically, findings from our lab and others in the norovirus field have defined opposing roles for bile acids in regulating norovirus infections: Host-derived bile acids promote human and murine norovirus infections in vitro through a range of mechanisms whereas microbiota-derived bile acids inhibit infection in adult mice via amplification of host antiviral immunity^14,15,22,23^. Considering the opposing proviral and antiviral effects of host-derived and microbiota-derived bile acids, respectively, on norovirus infection and the immaturity of gut microbiota and bile acid metabolism in neonates^26–28^, we hypothesized that the microbiota-regulated balance of the bile acid pool may be a key determinant of the pathogenic outcome of a norovirus infection. Our findings lend strong support to this concept: (i) the dominance of the neonatal bile acid pool by host-derived bile acids was associated with susceptibility to norovirus-induced disease; (ii) early postnatal microbiota maturation via reconstitution with bile acid-biotransforming bacteria protected from disease; and (iii) microbiota-derived bile acids, and specifically TGR5 ligands, protected neonates from disease whereas host-derived bile acids exacerbated disease. Importantly, bile acid metabolism is also immature in human newborns^21,26–29,64^ and human norovirus infections are enhanced by host-derived bile acids in vitro^22,65^ so it is entirely plausible that metabolic immaturity underlies neonatal susceptibility to severe norovirus disease in people as it does in mice. Moreover, microbiota-derived bile acids exert antiviral activity against a wide array of other viruses^13,24,25,66–68^, many of which also cause more severe disease in neonates than in adults. This raises the possibility that the paucity of microbiota-derived bile acids in neonatal hosts represents a common cause of neonatal vulnerability to viruses. More broadly, metabolites are central determinants of the host-pathogen interaction in many, if not most, infections^13,69^ so immaturity of other metabolic pathways in the developing offspring could underlie susceptibility to many types of infectious organisms.

Remarkably, we also discovered that maternal bile acids delivered to neonates via breastfeeding play a central role in determining neonatal susceptibility to norovirus diarrhea. While several previous studies reported detection of bile acids in human and rodent breast milk^30,31^, the source of these metabolites and their biological significance has been largely unexplored. Our results represent the most comprehensive evaluation of the breast milk bile acid landscape to date. Furthermore, they suggest that entero-mammary trafficking is the major source of bile acids in the mammary gland since gut microbiota-metabolized bile acids were present in breast milk and the composition of the breast milk bile acid pool was markedly altered by deficiency of the main intestinal bile acid transporter ASBT and by oral bile acid sequestration. In both human and mouse breast milk, the most abundant bile acids detected were host-derived conjugated primary species. These bile acids drove norovirus disease in breastfeeding mice: Pups fostered by dams lacking conjugated primary bile acids in their breast milk were entirely protected from disease and disease susceptibility was restored by oral administration of a host-derived bile acid to the neonate. While maternal bile acid delivery during breastfeeding undoubtedly plays beneficial roles in the developing offspring, our results reveal that it has the unintended negative consequence of heightening disease severity during an enteric virus infection.

Finally, our results in a small animal model of norovirus diarrhea provide support for novel infant-specific therapeutic strategies based on the finding that bile acid metabolism determines host susceptibility to norovirus disease. These include driving early maturation of the microbiota to accelerate colonization with bile acid-biotransforming bacterial species, activating key bile acid receptors like TGR5 to potentiate antiviral immune responses, and targeting entero-mammary trafficking of bile acids in the lactating mother.

## EXPERIMENTAL METHODS

### Viruses

BV2 and HEK293T cells were maintained in Dulbecco’s modified Eagle medium supplemented with 10% fetal bovine serum, 100 U penicillin/mL, and 100 μg/mL streptomycin. Cells tested negative for mycoplasma. A MNV WU23 (GenBank accession number EU004668.1; A4826C and G4828A mutations added) infectious clone^33^ was used to generate virus stocks. In brief, 10^6^ HEK293T cells were transfected with 5 µg of endotoxin-free infectious clone using Lipofectamine 3000. Cells were frozen after 30 h, lysed by freeze-thaw, and lysates applied to BV2 cells. BV2 lysates were frozen when cultures displayed 90% cytopathic effect and then freeze-thawed twice, cell debris removed by low-speed centrifugation, followed by purification of virus through a 25% sucrose cushion. Virus was dissolved in phosphate-buffered saline (PBS). Virus stocks were titered by a standard TCID_50_ assay as described previously^70^. Stocks were sequenced to confirm no mutations arose during generation. A mock inoculum stock was prepared in the same manner using BV2 lysate from uninfected cultures.

### Mice

Specific pathogen-free (SPF) mice used in this study were bred and housed in animal facilities at the University of Florida under strict MNV-free conditions. Germ-free C57BL/6 mice used in this study were originally obtained from Taconic Bioscience and were bred and housed in animal facilities at the University of Illinois at Chicago under sterile conditions in isolators and provided with sterile water and sterile standard pellet diet ad libitum on a 12 h light-dark cycle at 22C. Standard microbiological methods were used to check fecal samples weekly for yeast, molds, and aerobic and anaerobic bacteria to confirm the germ-free conditions in the isolators. All animal experiments were performed in strict accordance with federal and university guidelines and approved by the Institutional Animal Care and Use Committee at the University of Florida (study numbers 202110473 and 202200000065) and University of Illinois at Chicago (study number ACC#22-110). The conditions in animal rooms used in this study fall within the standards set by the “Guide for the Care and Use of Laboratory Animals.” C57BL/6J mice (Jackson no. 000664, referred to as B6), C57BL/6J-*Gpbar1^-/-^* mice^71^ (kindly provided by Wendong Huang, Beckman Research Institute of City of Hope; referred to as *Tgr5^-/-^*) and C57BL/6NJ-*Slc10a^-/-^*mice^72^ (kindly provided by Paul Dawson, Emory University; referred to as *Asbt^-/-^*) mice were used.

### Neonatal virus infections and disease measurements

For all neonatal mouse infections except where otherwise indicated, postnatal day 3 (P3) neonatal mice were inoculated with 1 x 10^7^ TCID50 units of WU23 unless otherwise indicated or mock inoculum using intragastric (i.g.) inoculation, as previously published^33^. In mice P7 or older, oral gavage was used for inoculations instead since the milk spot was no longer visible^32^. For assessing diarrhea in time course experiments, the abdomen of each pup was palpated to induce defecation at indicated time points. Fecal condition was assessed based on color and consistency according to a 6-point scale: 0, no defecation; 1, firm, orange, does not smear; 2, pasty, orange or mixed color, does not smear; 3, orange or yellow, semi-liquid and smears; 4, yellow, liquid, and smears; 5, green-yellow, non-viscous liquid. Any pup that scored a 3-5 was considered to have diarrhea for the purpose of calculating incidence. Scores of 0 were excluded from analysis since it was not possible to assess consistency. For assessing diarrhea at individual time points, the consistency of colon contents was scored after sacrificing the mouse utilizing the same scoring system. We have determined that fecal and colon content scores are highly concordant and can be used interchangeably^34^. For measuring intestinal permeability, neonates were separated from dams 3 h prior to the end of the experiment, weighed, and injected i.g. with 40 mg/kg of fluorescein isothiocyanate-dextran 10kDa (FD10; Sigma-Aldrich, FD10S) in sterile double-distilled water (ddH2O). At 3 h post-FD10 administration, neonates were scored for disease and sera was collected in BD microtainer blood collection tubes. The sera were centrifuged for 5 min at 4C, diluted 40x in sterile ddH2O, and plated in triplicate in a 96-well plate. Samples were read for absorbance at an excitation of 490 nm and emission of 520 nm on a Spectra Max2 microplate reader. Values were calculated using a standard curve.

### In vivo microbiota and bile acid manipulations

For microbiota depletions, 90 mg/kg of vancomycin (Fisher Scientific), ampicillin (Sigma-Aldrich), and gentamicin (Sigma-Aldrich) were diluted in sterile 1X PBS for a total of 40 μL per neonate and delivered i.g. to neonates 1 d prior to infection and 1 dpi. Control mice received PBS alone. For P7 and P10 mice used in age restriction studies, the antibiotic cocktail (ABX) was administered by oral gavage. In all microbiota depletion studies, the same ABX was added to the dam’s drinking water at a final concentration of 1 mg/mL of each antibiotic 1 d prior to neonatal ABX administration. To confirm efficient depletions, fecal samples from neonates and dams were collected at the time of infection, homogenized, plated on brain-heart infusion agar with 10% sheep blood, and cultured under anaerobic or aerobic conditions at 37°C for 2 d. Bacterial depletion was confirmed by an absence of growth. For bacterial reconstitutions, the *Clostridium scindens* strain VPI 13733 (ATCC 35704) was cultured in degassed tryptic soy medium with 5% defibrinated sheep blood (ATCC medium 260) for 3 d at 37C in an anaerobic chamber (Coy Lab Products Vinyl Anaerobic Chamber Type B) kept at 5% CO2 in hydrogen-balanced nitrogen. Neonatal mice were administered ABX at P2 and inoculated with 10^6^ colony forming units (CFU) of *C. scindens* at P3. Control mice were treated with ABX only or ABX plus ATCC medium 260. For caffeic phenyl ester (CAPE) treatment, mice were treated with 75 mg/kg CAPE (MedChem Express) or DMSO in corn oil as vehicle control at P4 and P6. For bile acid supplementations of neonatal mice, mice were ABX-treated at P2, P4, and P6 and were administered 70 μg/g of deoxycholic acid (DCA), taurocholic acid (TCA), or vehicle control at P3, P4, and P5 i.g. Corn oil was used as a vehicle control for DCA and PBS was used for TCA. Mice were infected with WU23 at P5.5. For TGR5 agonist treatment, mice were administered 60 mg/kg of INT-777 (MedChem Express) in corn oil in 20 µL via i.g. on P2, P3, P4, and P6 and infected on P5. For bile acid sequestration in lactating dams, dams were fed chow supplemented with 2% cholestyramine (Fisher) or control chow starting at P0 and maintained throughout the experiment. Chow was replaced daily.

### Bile acid profiling

For mouse fecal bile acid profiling, stool was collected from naïve P4 and P5 neonatal mice and pooled, and from naïve 8-week-old adult mice. Each sample was mixed with 20 μL of acetonitrile per mg of raw material. The samples were then homogenized on a MM 400 mixer mill at 30 Hz for 3 min, followed by sonication for 3 min in an ice-water bath. The samples were centrifuged at 21,000 g for 10 min. 20 μL of the clear supernatant of each sample was mixed with 180 μL of the internal standard solution. For mouse breast milk bile acid profiling, lactating dams were milked daily on P8-P11 and pooled. On the day of milking, litters were separated from dams for 2 h and dams were given 2 international unit (IU)/kg of oxytocin diluted in saline (100 μl total volume) intraperitoneally (i.p.) and anesthetized using isoflurane. Eye lubricant was used for the duration of the procedure. Once anesthetized, milk was expressed from each mammary gland, collected using a micropipette, and frozen. Anesthesia time never exceeded 25 min total time. When the procedure was complete, the dams were monitored to ensure no side effects. An existing set of 24 human breast milk samples collected from women at various times 1-15 months following birth^73^ were used for bile acid profiling. The human study was approved by the Institutional Review Boards of the National Autonomous University of Nicaragua, León (UNAN-León, Acta Number 45, 2017) and the University of North Carolina at Chapel Hill (Study Number: 16-2079). 20 μL of each sample was mixed with 80 μL of acetonitrile. After vortexing, sonication for 5 min in an ice-water bath, and centrifugation at 21,000 g for 15 min, 80 μL of the clear supernatant was mixed with 40 μL of the internal standard solution and 880 μL of water. The mixture was loaded onto a reversed-phase solid-phase extraction cartridge (60mg/1mL). After sample loading under a positive pressure, the cartridge was washed with 2 mL of water. Bile acids were eluted with 1 mL of methanol under the positive pressure. The collected fraction was dried in a nitrogen gas evaporator. The residue was reconstituted in 40 μL of 50% acetonitrile. 10 μL aliquots of each fecal and milk sample solutions and each of the calibration solutions were then injected into an Agilent 1290 UHPLC system coupled to an Agilent 6495B QQQ mass spectrometer. The MS instrument was operated in the multiple-reaction monitoring (MRM) mode with negative ion detection. A Waters C18 column (2.1*150 mm, 1.7 μm) was used for LC separation and the mobile phase was 0.01% formic acid in water and in acetonitrile for binary-solvent gradient elution. A mixed solution of all the targeted bile acids at 10 μM of each compound was prepared in an internal standard solution of 14 deuterium-labeled bile acids in 50% acetonitrile. This solution was further diluted step by step to have 10 calibration solutions. Linear-regression calibration curves of individual bile acids were constructed with the data acquired from injection of the serially diluted calibration solutions. Concentrations of bile acids detected in the samples were calculated by interpolating the calibration curves of individual bile acids with the analyte-to-internal standard peak area ratios measured form injection of the sample solutions. Bile acid profiling was performed by Creative Proteomics.

### RNA extraction and quantitative RT-PCR

RNA was isolated from the indicated segment of ileal tissue from neonatal or adult mice using the RNeasy Plus Kit (Qiagen), treated with Turbo DNase (Invitrogen), and 1 μg RNA was used for cDNA synthesis with the High-Capacity cDNA Reverse Transcription Kit (ThermoFisher). Quantitative PCR (qPCR) was performed using the SYBR Green Master Mix (ThermoFisher) using primers specific to the ASBT gene (F: 5’ttgcctcttcgtctacacc3’; R: 5’ccaaaggaaacaggaataacaag3’), IFN-β gene (F: 5’gcagctgaatggaaagatca3’; R: 5’gtggagagcagttgaggaca3’), and GAPDH housekeeping gene (F: 5’catggccttccgtgttccta3’; R: 5’cctgcttcaccaccttcttgat3’). ASBT and IFN-β expression was normalized to GAPDH and all samples were analyzed with technical triplicates.

### RNAscope-based in situ hybridization (ISH)

RNAscope ISH assays were performed as previously described^74^. In brief, small intestine sections were fixed in 10% buffered formalin for 16 h and then transferred to PBS. Tissues were paraffin-embedded within 24 h and sections of 4-µm thickness applied to glass slides. Tissues were deparaffinized by heating at 60C for 30 min followed by xylene treatment and dehydration. Sections were then hybridized with a custom-designed probe targeting SLC10A2 (ACDBio 1215641) for 2 h at 40C followed by probe amplification and detection using the ACDBio RNAscope RED detection kit. Positive (UBC) and negative (DapB) control probes were stained in parallel for all experiments.

### Immunofluorescence assays

Small intestine sections prepared as described above were deparaffinized by xylene treatment and rehydrated, antigen retrieval was performed by incubating slides in sodium citrate buffer (8.2 mM sodium citrate, 1.8 mM citric acid) for 15 min at high pressure in a pressure cooker, sections were blocked for 1 h at room temperature in serum-free blocking buffer (Agilent Technologies, X090930-2), and then incubated with polyclonal rabbit antibody to ASBT (kindly provided by Paul Dawson, Emory University) at a 1:400 dilution overnight at 4C. Negative controls were incubated with serum-free blocking buffer. All slides were subsequently incubated with 1:5000 dilution Alexa Fluor 647-conjugated anti-rabbit antibody (Invitrogen). Nuclei were visualized with DAPI (4′,6′-diamidino-2-phenylindole)-containing Fluoroshield mounting medium (Sigma-Aldrich). A Nikon A1RMPsi laser scanning confocal microscope was used for imaging. Images were analyzed using ImageJ Fiji and NIS Elements Viewer software.

### Statistical analysis

All data were analyzed with GraphPad Prism software. *P* values were determined using one-way or two-way ANOVA with corrections for multiple comparisons. Error bars denote standard errors of mean in all figures.

## Acknowledgements

We thank the University of Illinois at Chicago Gnotobiotic Mouse Facility for maintaining the germ-free colony used in our studies.

## Funding

This work was supported by the National Institutes of Health (NIH) grants R01AI162970 (S.M.K.), R01AI141478 (S.M.K.), T32AI007110 (A.M.P. and J.M.A.), F30AI172364 (A.M.P.), F30AI154834 (E.W.H.), R01DK123826 (X.D.T.), R01DK129960 (X.D.T.), R01DK047987 (P.A.D.), and US Department of Veterans Affairs Merit Review Award I01BX001690 (X.D.T.). Imaging done on the Nikon A1RMPsi system was provided by the NIH grant 1S10OD020026.

## Competing interests

We declare no competing interests.

## Data and materials availability

The data from this study are tabulated in the main paper and supplementary materials. All reagents are available from S.M.K. under a material transfer agreement with University of Florida.

## Notes

### Competing Interest Statement

The authors have declared no competing interest.

## References

1. Koo HL, Neill FH, Estes MK, Munoz FM, Cameron A, Dupont HL, Atmar RL. Noroviruses: The Most Common Pediatric Viral Enteric Pathogen at a Large University Hospital After Introduction of Rotavirus Vaccination. J Pediatric Infect Dis Soc. 2013 Mar;2(1):57–60. PMID: 23687584

2. Patel MM. Systematic Literature Review of Role of Noroviruses in Sporadic Gastroenteritis. Emerg Infect Dis. 2008 Aug;14(8):1224–1231.

3. Ahmed SM, Hall AJ, Robinson AE, Verhoef L, Premkumar P, Parashar UD, Koopmans M, Lopman BA. Global prevalence of norovirus in cases of gastroenteritis: a systematic review and meta-analysis. The Lancet Infectious Diseases. 2014 Aug;14(8):725–730.

4. Alard P. Gut microbiota, immunity, and disease: a complex relationship. Front Microbio. 2011;2:180.

5. Thaiss CA, Zmora N, Levy M, Elinav E. The microbiome and innate immunity. Nature. 2016 Jul 7;535(7610):65–74.

6. Abt MC, Artis D. The intestinal microbiota in health and disease: the influence of microbial products on immune cell homeostasis. Current Opinion in Gastroenterology. 2009 Nov;25(6):496–502.

7. Jones MK, Watanabe M, Zhu S, Graves CL, Keyes LR, Grau KR, Gonzalez-Hernandez MB, Iovine NM, Wobus CE, Vinjé J, Tibbetts SA, Wallet SM, Karst SM. Enteric bacteria promote human and mouse norovirus infection of B cells. Science. 2014 Nov 7;346(6210):755–759. PMCID: PMC4401463

8. Kernbauer E, Ding Y, Cadwell K. An enteric virus can replace the beneficial function of commensal bacteria. Nature. 2014 Dec 4;516(7529):94–98.

9. Baldridge MT, Nice TJ, McCune BT, Yokoyama CC, Kambal A, Wheadon M, Diamond MS, Ivanova Y, Artyomov M, Virgin HW. Commensal microbes and interferon-λ determine persistence of enteric murine norovirus infection. Science. 2015;347(6219):266–9. PMID: 25431490

10. Kuss SK, Best GT, Etheredge CA, Pruijssers AJ, Frierson JM, Hooper LV, Dermody TS, Pfeiffer JK. Intestinal Microbiota Promote Enteric Virus Replication and Systemic Pathogenesis. Science. 2011 Oct 14;334(6053):249–252.

11. Kane M, Case LK, Kopaskie K, Kozlova A, MacDearmid C, Chervonsky AV, Golovkina TV. Successful Transmission of a Retrovirus Depends on the Commensal Microbiota. Science. 2011 Oct 14;334(6053):245–249.

12. Uchiyama R, Chassaing B, Zhang B, Gewirtz AT. Antibiotic Treatment Suppresses Rotavirus Infection and Enhances Specific Humoral Immunity. J Infect Dis. 2014 Jul 15;210(2):171–182. PMID: 24436449

13. Alwin A, Karst SM. The influence of microbiota-derived metabolites on viral infections. Current Opinion in Virology. 2021 Aug 1;49:151–156.

14. Nelson CA, Wilen CB, Dai YN, Orchard RC, Kim AS, Stegeman RA, Hsieh LL, Smith TJ, Virgin HW, Fremont DH. Structural basis for murine norovirus engagement of bile acids and the CD300lf receptor. Proc Natl Acad Sci USA. 2018 25;115(39):E9201–E9210. PMCID: PMC6166816

15. Sherman MB, Williams AN, Smith HQ, Nelson C, Wilen CB, Fremont DH, Virgin HW, Smith TJ. Bile Salts Alter the Mouse Norovirus Capsid Conformation: Possible Implications for Cell Attachment and Immune Evasion. Journal of Virology. American Society for Microbiology Journals; 2019 Oct 1;93(19):e00970–19. PMID: 31341042

16. Creutznacher R, Schulze E, Wallmann G, Peters T, Stein M, Mallagaray A. Chemical-Shift Perturbations Reflect Bile Acid Binding to Norovirus Coat Protein: Recognition Comes in Different Flavors. Chembiochem. 2020 Apr 1;21(7):1007–1021. PMCID: PMC7186840

17. Kilic T, Koromyslova A, Hansman GS. Structural Basis for Human Norovirus Capsid Binding to Bile Acids. J Virol. 2019 Jan 4;93(2):e01581–18. PMCID: PMC6321941

18. Kong F, Saif LJ, Wang Q. Roles of bile acids in enteric virus replication. Animal Diseases. 2021 Apr 23;1(1):2.

19. Bordon Y. Bacterial metabolites shape neonatal immune system. Nature Reviews Immunology. Nature Publishing Group; 2019 Sep;19(9):537–537.

20. Kim CH. Immune regulation by microbiome metabolites. Immunology. 2018;154(2):220–229. PMCID: PMC5980225

21. Craddock AL, Love MW, Daniel RW, Kirby LC, Walters HC, Wong MH, Dawson PA. Expression and transport properties of the human ileal and renal sodium-dependent bile acid transporter. Am J Physiol. 1998 Jan;274(1):G157–169. PMID: 9458785

22. Murakami K, Tenge VR, Karandikar UC, Lin SC, Ramani S, Ettayebi K, Crawford SE, Zeng XL, Neill FH, Ayyar BV, Katayama K, Graham DY, Bieberich E, Atmar RL, Estes MK. Bile acids and ceramide overcome the entry restriction for GII.3 human norovirus replication in human intestinal enteroids. Proc Natl Acad Sci U S A. 2020 Jan 21;117(3):1700–1710. PMCID: PMC6983410

23. Grau KR, Zhu S, Peterson ST, Helm EW, Philip D, Phillips M, Hernandez A, Turula H, Frasse P, Graziano VR, Wilen CB, Wobus CE, Baldridge MT, Karst SM. The intestinal regionalization of acute norovirus infection is regulated by the microbiota via bile acid-mediated priming of type III interferon. Nat Microbiol. Nature Publishing Group; 2020 Jan;5(1):84–92.

24. Winkler ES, Shrihari S, Hykes BL, Handley SA, Andhey PS, Huang YJS, Swain A, Droit L, Chebrolu KK, Mack M, Vanlandingham DL, Thackray LB, Cella M, Colonna M, Artyomov MN, Stappenbeck TS, Diamond MS. The Intestinal Microbiome Restricts Alphavirus Infection and Dissemination through a Bile Acid-Type I IFN Signaling Axis. Cell. 2020 Aug 20;182(4):901–918.e18.

25. Nagai M, Moriyama M, Ishii C, Mori H, Watanabe H, Nakahara T, Yamada T, Ishikawa D, Ishikawa T, Hirayama A, Kimura I, Nagahara A, Naito T, Fukuda S, Ichinohe T. High body temperature increases gut microbiota-dependent host resistance to influenza A virus and SARS-CoV-2 infection. Nat Commun. 2023 Jun 30;14:3863. PMCID: PMC10313692

26. Obinata K, Nittono H, Yabuta K, Mahara R, Tohma M. Metabolism of Fetal Bile Acids in Healthy Neonates. Pediatr Res. Nature Publishing Group; 1989 Sep;26(3):273–273.

27. van Best N, Rolle-Kampczyk U, Schaap FG, Basic M, Olde Damink SWM, Bleich A, Savelkoul PHM, von Bergen M, Penders J, Hornef MW. Bile acids drive the newborn’s gut microbiota maturation. Nat Commun. 2020 Jul 23;11:3692. PMCID: PMC7378201

28. Tanaka M, Sanefuji M, Morokuma S, Yoden M, Momoda R, Sonomoto K, Ogawa M, Kato K, Nakayama J. The association between gut microbiota development and maturation of intestinal bile acid metabolism in the first 3 y of healthy Japanese infants. Gut Microbes. 2019 Sep 24;11(2):205–216. PMCID: PMC7053967

29. Shneider BL, Setchell KDR, Crossman MW. Fetal and Neonatal Expression of the Apical Sodium-Dependent Bile Acid Transporter in the Rat Ileum and Kidney. Pediatr Res. Nature Publishing Group; 1997 Aug;42(2):189–194.

30. Forsyth JS, Ross PE, Bouchier IA. Bile salts in breast milk. Eur J Pediatr. 1983 Apr;140(2):126–127. PMID: 6884389

31. Blazquez AMG, Macias RIR, Cives-Losada C, de la Iglesia A, Marin JJG, Monte MJ. Lactation during cholestasis: Role of ABC proteins in bile acid traffic across the mammary gland. Sci Rep. 2017 Aug 7;7:7475. PMCID: PMC5547141

32. Roth AN, Helm EW, Mirabelli C, Kirsche E, Smith JC, Eurell LB, Ghosh S, Altan-Bonnet N, Wobus CE, Karst SM. Norovirus infection causes acute self-resolving diarrhea in wild-type neonatal mice. Nature Communications. Nature Publishing Group; 2020 Jun 11;11(1):2968.

33. Peiper AM, Helm EW, Nguyen Q, Phillips M, Williams CG, Shah D, Tatum S, Iyer N, Grodzki M, Eurell LB, Nasir A, Baldridge MT, Karst SM. Infection of neonatal mice with the murine norovirus strain WU23 is a robust model to study norovirus pathogenesis. Lab Animal. 2023;52(6):119–129.

34. Helm EW, Peiper AM, Phillips M, Williams CG, Sherman MB, Kelley T, Smith HQ, Jacobs SO, Shah D, Tatum SM, Iyer N, Grodzki M, Morales Aparicio JC, Kennedy EA, Manzi MS, Baldridge MT, Smith TJ, Karst SM. Environmentally-triggered contraction of the norovirus virion determines diarrheagenic potential. Fronties in Immunology. 2022;13:1043746.

35. Clark RH, Bloom BT, Spitzer AR, Gerstmann DR. Reported medication use in the neonatal intensive care unit: data from a large national data set. Pediatrics. 2006 Jun;117(6):1979–1987. PMID: 16740839

36. Gopinath S, Kim MV, Rakib T, Wong PW, van Zandt M, Barry NA, Kaisho T, Goodman AL, Iwasaki A. Topical application of aminoglycoside antibiotics enhances host resistance to viral infections in a microbiota-independent manner. Nat Microbiol. 2018;3(5):611–621. PMCID: PMC5918160

37. Acevedo MAW, Pfeiffer JK. Microbiota-Independent Antiviral Effects of Antibiotics on Poliovirus and Coxsackievirus. Virology. 2020 Jul;546:20–24. PMCID: PMC7253499

38. Grunicke H, Pushendorf B, Werchau H. Mechanism of action of distamycin A and other antibiotics with antiviral activity. Rev Physiol Biochem Pharmacol. 1976;75:69–96. PMID: 59940

39. Zeng S, Meng X, Huang Q, Lei N, Zeng L, Jiang X, Guo X. Spiramycin and azithromycin, safe for administration to children, exert antiviral activity against enterovirus A71 in vitro and in vivo. Int J Antimicrob Agents. 2019 Apr;53(4):362–369. PMID: 30599241

40. Bawage SS, Tiwari PM, Pillai S, Dennis VA, Singh SR. Antibiotic Minocycline Prevents Respiratory Syncytial Virus Infection. Viruses. 2019 Aug 11;11(8):E739. PMCID: PMC6723987

41. Kang DJ, Ridlon JM, Moore DR, Barnes S, Hylemon PB. Clostridium scindens baiCD and baiH genes encode stereo-specific 7alpha/7beta-hydroxy-3-oxo-delta4-cholenoic acid oxidoreductases. Biochim Biophys Acta. 2008 Feb;1781(1–2):16–25. PMCID: PMC2275164

42. Bourgin M, Kriaa A, Mkaouar H, Mariaule V, Jablaoui A, Maguin E, Rhimi M. Bile Salt Hydrolases: At the Crossroads of Microbiota and Human Health. Microorganisms. Multidisciplinary Digital Publishing Institute; 2021 Jun;9(6):1122.

43. Smith K, Zeng X, Lin J. Discovery of Bile Salt Hydrolase Inhibitors Using an Efficient High-Throughput Screening System. PLOS ONE. Public Library of Science; 2014 Jan 14;9(1):e85344.

44. Bertolini A, Fiorotto R, Strazzabosco M. Bile acids and their receptors: modulators and therapeutic targets in liver inflammation. Semin Immunopathol. 2022 Jul;44(4):547–564. PMCID: PMC9256560

45. Jia W, Xie G, Jia W. Bile acid–microbiota crosstalk in gastrointestinal inflammation and carcinogenesis. Nature Reviews Gastroenterology and Hepatology. 2017 Oct 11;nrgastro.2017.119.

46. Xiong Q, Huang H, Wang N, Chen R, Chen N, Han H, Wang Q, Siwko S, Liu M, Qian M, Du B. Metabolite-Sensing G Protein Coupled Receptor TGR5 Protects Host From Viral Infection Through Amplifying Type I Interferon Responses. Front Immunol. 2018;9:2289. PMCID: PMC6176213

47. Hu MM, He WR, Gao P, Yang Q, He K, Cao LB, Li S, Feng YQ, Shu HB. Virus-induced accumulation of intracellular bile acids activates the TGR5-β-arrestin-SRC axis to enable innate antiviral immunity. Cell Research. Nature Publishing Group; 2019 Mar;29(3):193–205.

48. Xiong F, Cao L, Wu XM, Chang MX. The function of zebrafish gpbar1 in antiviral response and lipid metabolism. Developmental & Comparative Immunology. 2021 Mar 1;116:103955.

49. Pellicciari R, Gioiello A, Macchiarulo A, Thomas C, Rosatelli E, Natalini B, Sardella R, Pruzanski M, Roda A, Pastorini E, Schoonjans K, Auwerx J. Discovery of 6α-Ethyl-23(S)-methylcholic Acid (S-EMCA, INT-777) as a Potent and Selective Agonist for the TGR5 Receptor, a Novel Target for Diabesity. J Med Chem. American Chemical Society; 2009 Dec 24;52(24):7958–7961.

50. Hughes KR, Schofield Z, Dalby MJ, Caim S, Chalklen L, Bernuzzi F, Alcon-Giner C, Le Gall G, Watson AJM, Hall LJ. The early life microbiota protects neonatal mice from pathological small intestinal epithelial cell shedding. The FASEB Journal. 2020;34(5):7075–7088.

51. Ferrebee CB, Li J, Haywood J, Pachura K, Robinson BS, Hinrichs BH, Jones RM, Rao A, Dawson PA. Organic Solute Transporter α-β Protects Ileal Enterocytes From Bile Acid–Induced Injury. Cell Mol Gastroenterol Hepatol. 2018 Jan 12;5(4):499–522. PMCID: PMC6009794

52. Cui JY, Aleksunes LM, Tanaka Y, Fu ZD, Guo Y, Guo GL, Lu H, Zhong X bo, Klaassen CD. Bile acids via FXR initiate the expression of major transporters involved in the enterohepatic circulation of bile acids in newborn mice. Am J Physiol Gastrointest Liver Physiol. 2012 May 1;302(9):G979–G996. PMCID: PMC3362079

53. Rodríguez JM, Fernández L, Verhasselt V. The Gut‒Breast Axis: Programming Health for Life. Nutrients. 2021 Feb 12;13(2):606. PMCID: PMC7917897

54. Christie DM, Dawson PA, Thevananther S, Shneider BL. Comparative analysis of the ontogeny of a sodium-dependent bile acid transporter in rat kidney and ileum. Am J Physiol. 1996 Aug;271(2 Pt 1):G377-385. PMID: 8770054

55. Chen F, Shyu AB, Shneider BL. Hu antigen R and tristetraprolin: Counter-regulators of rat apical sodium-dependent bile acid transporter by way of effects on messenger RNA stability. Hepatology. 2011;54(4):1371–1378.

56. Hwang ST, Henning SJ. Ontogenic regulation of components of ileal bile acid absorption. Exp Biol Med (Maywood). 2001 Jul;226(7):674–680. PMID: 11444103

57. Schuck-Phan A, Phan T, Dawson P, Dial E, Bell C, Liu Y, Rhoads J, Lichtenberger L. Formula Feeding Predisposes Gut to NSAID-Induced Small Intestinal Injury. Clin Exp Pharmacol. 2016;6(6):222. PMCID: PMC6764459

58. Xiao Y, Yan W, Zhou K, Cao Y, Cai W. Glucocorticoid treatment alters systemic bile acid homeostasis by regulating the biosynthesis and transport of bile salts. Digestive and Liver Disease. 2016 Jul 1;48(7):771–779.

59. Halpern MD, Holubec H, Saunders TA, Dvorak K, Clark JA, Doelle SM, Ballatori N, Dvorak B. Bile Acids Induce Ileal Damage During Experimental Necrotizing Enterocolitis. Gastroenterology. 2006 Feb;130(2):359–372. PMCID: PMC3417808

60. Halpern MD, Weitkamp JH, Mount Patrick SK, Dobrenen HJ, Khailova L, Correa H, Dvorak B. Apical sodium-dependent bile acid transporter upregulation is associated with necrotizing enterocolitis. Am J Physiol Gastrointest Liver Physiol. 2010 Sep;299(3):G623–G631. PMCID: PMC2950692

61. Cannon JL, Lopman BA, Payne DC, Vinjé J. Birth Cohort Studies Assessing Norovirus Infection and Immunity in Young Children: A Review. Clin Infect Dis. 2019 Jul 2;69(2):357–365.

62. Menon VK, George S, Sarkar R, Giri S, Samuel P, Vivek R, Saravanabavan A, Liakath FB, Ramani S, Iturriza-Gomara M, Gray JJ, Brown DW, Estes MK, Kang G. Norovirus Gastroenteritis in a Birth Cohort in Southern India. PLOS ONE. 2016 Jun 10;11(6):e0157007.

63. Rouhani S, Peñataro Yori P, Paredes Olortegui M, Siguas Salas M, Rengifo Trigoso D, Mondal D, Bodhidatta L, Platts-Mills J, Samie A, Kabir F, Lima A, Babji S, Mason CJ, Kalam A, Bessong P, Ahmed T, Mduma E, Bhutta ZA, Lima I, Ramdass R, Lang D, George A, Zaidi AKM, Kang G, Houpt E, Kosek MN, Etiology, Risk Factors, and Interactions of Enteric Infections and Malnutrition and the Consequences for Child Health and Development Project (MAL-ED) Network Investigators. Norovirus Infection and Acquired Immunity in 8 Countries: Results From the MAL-ED Study. Clin Infect Dis. 2016 15;62(10):1210–1217. PMCID: PMC4845786

64. Lynch LE, Hair AB, Soni KG, Yang H, Gollins LA, Narvaez-Rivas M, Setchell KDR, Preidis GA. Cholestasis impairs gut microbiota development and bile salt hydrolase activity in preterm neonates. Gut Microbes. 15(1):2183690. PMCID: PMC9980517

65. Ettayebi K, Crawford SE, Murakami K, Broughman JR, Karandikar U, Tenge VR, Neill FH, Blutt SE, Zeng XL, Qu L, Kou B, Opekun AR, Burrin D, Graham DY, Ramani S, Atmar RL, Estes MK. Replication of human noroviruses in stem cell–derived human enteroids. Science. 2016 Aug 25;353(6306):1387–1393. PMID: 27562956

66. Xiong Q, Huang H, Wang N, Chen R, Chen N, Han H, Wang Q, Siwko S, Liu M, Qian M, Du B. Metabolite-Sensing G Protein Coupled Receptor TGR5 Protects Host From Viral Infection Through Amplifying Type I Interferon Responses. Front Immunol. 2018;9:2289. PMCID: PMC6176213

67. Hu MM, He WR, Gao P, Yang Q, He K, Cao LB, Li S, Feng YQ, Shu HB. Virus-induced accumulation of intracellular bile acids activates the TGR5-β-arrestin-SRC axis to enable innate antiviral immunity. Cell Res. 2019 Mar;29(3):193–205. PMCID: PMC6460433

68. Schupp AK, Trilling M, Rattay S, Le-Trilling VTK, Haselow K, Stindt J, Zimmermann A, Häussinger D, Hengel H, Graf D. Bile Acids Act as Soluble Host Restriction Factors Limiting Cytomegalovirus Replication in Hepatocytes. J Virol. 2016 Jul 11;90(15):6686–6698. PMCID: PMC4944301

69. Li Z, Quan G, Jiang X, Yang Y, Ding X, Zhang D, Wang X, Hardwidge PR, Ren W, Zhu G. Effects of Metabolites Derived From Gut Microbiota and Hosts on Pathogens. Front Cell Infect Microbiol. 2018 Sep 14;8:314. PMCID: PMC6152485

70. Thackray LB, Wobus CE, Chachu KA, Liu B, Alegre ER, Henderson KS, Kelley ST, Virgin HW. Murine Noroviruses Comprising a Single Genogroup Exhibit Biological Diversity despite Limited Sequence Divergence. J Virol. 2007 Oct 1;81(19):10460–10473.

71. Vassileva G, Golovko A, Markowitz L, Abbondanzo SJ, Zeng M, Yang S, Hoos L, Tetzloff G, Levitan D, Murgolo NJ, Keane K, Davis HR, Hedrick J, Gustafson EL. Targeted deletion of Gpbar1 protects mice from cholesterol gallstone formation. Biochem J. 2006 Sep 15;398(3):423–430. PMCID: PMC1559456

72. Truong JK, Li J, Li Q, Pachura K, Rao A, Gumber S, Fuchs CD, Feranchak AP, Karpen SJ, Trauner M, Dawson PA. Active enterohepatic cycling is not required for the choleretic actions of 24-norUrsodeoxycholic acid in mice. JCI Insight. 8(6):e149360. PMCID: PMC10070106

73. Reyes Y, González F, Gutiérrez L, Blandón P, Centeno E, Zepeda O, Toval-Ruíz C, Lindesmith LC, Baric RS, Vielot N, Diez-Valcarce M, Vinjé J, Svensson L, Becker-Dreps S, Nordgren J, Bucardo F. Secretor Status Strongly Influences the Incidence of Symptomatic Norovirus Infection in a Genotype-Dependent Manner in a Nicaraguan Birth Cohort. J Infect Dis. 2022 Jan 5;225(1):105–115. PMCID: PMC8730499

74. Grau KR, Roth AN, Zhu S, Hernandez A, Colliou N, DiVita BB, Philip DT, Riffe C, Giasson B, Wallet SM, Mohamadzadeh M, Karst SM. The major targets of acute norovirus infection are immune cells in the gut-associated lymphoid tissue. Nature Microbiology. 2017 Dec;2(12):1586.

