## Supplemental Data for "Metabolic immaturity of newborns and breast milk bile acid metabolites are the central determinants of heightened neonatal vulnerability to norovirus diarrhea"

**Supplementary Figure 1. Positive correlation between fecal scores and intestinal permeability**

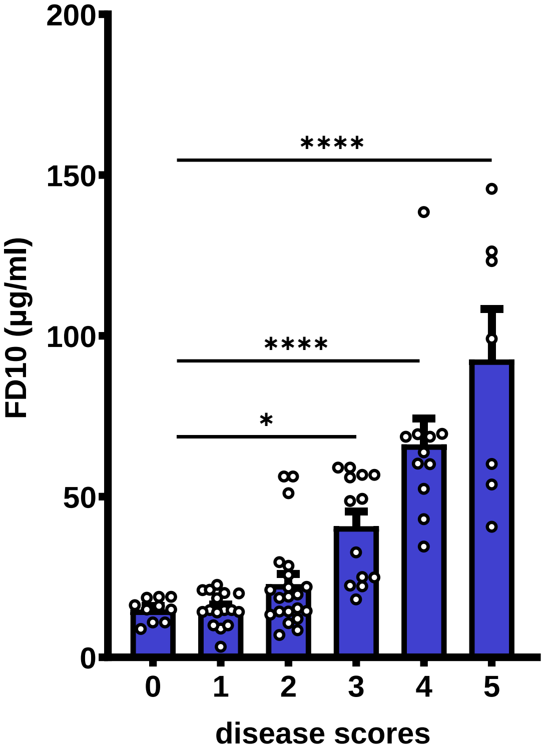


**Supplementary Figure 1**. **Intestinal permeability positively correlates with fecal scores during MNV infection.** Neonatal B6 mice (*n* = 78) were infected i.g. with 1 x $10^7$ $TCID_{50}$ units of WU23 or mock inoculum at P3, 40 mg/kg of FD10 was administered i.g. at 69 hpi, and the level of FD10 in serum was measured at 72 hpi. Disease severity was also assessed at 72 hpi by fecal scores. Fecal scores of 3 or higher indicate intestinal disease. FD10 measurements were plotted against the corresponding fecal score.

**Supplementary Figure 2. ASBT gene expression in neonatal and adult mice**

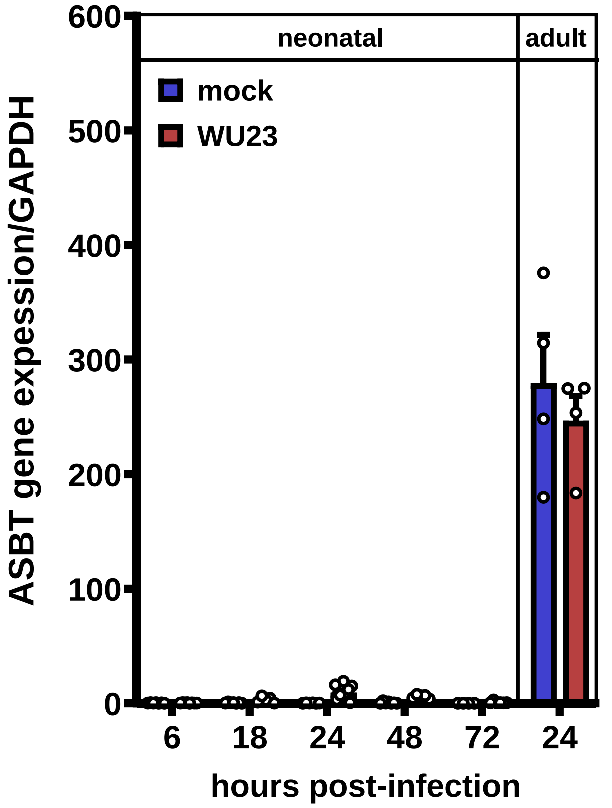


**Supplementary Figure 2. MNV infection does not prematurely induce expression of ASBT in neonatal mice.** Groups of P3 neonatal and 8-week-old adult B6 mice were infected with 1 x $10^7$ $TCID_{50}$ units of WU23 or mock inoculum. Portions of the neonatal distal small intestines were harvested either at 6 hpi (mock: *n* = 7; WU23: *n* = 8), 18 hpi (mock: *n* = 7; WU23: *n* = 5), 24 hpi (mock: *n* = 6; WU23: *n* = 12), 48 hpi (mock: *n* = 7; WU23: *n* = 5), or 72 hpi (mock: *n* = 4; WU23: *n* = 6), and adult tissue was harvested at 24 hpi (mock: *n* = 4; WU23: *n* = 4). RNA was extracted and qPCR was performed with primers specific to *ASBT* and normalized to the *GAPDH* housekeeping gene.
